# Fine-tuned method to extract high purified proteins from the seagrass *Halophila stipulacea* to be used for proteome analyses

**DOI:** 10.1101/2021.07.23.453549

**Authors:** Amalia Piro, Vasileia Anagnostopoulou, Eugenia T. Apostolaki, Silvia Mazzuca

## Abstract

The non-indigenous to the Mediterranean tropical seagrass *Halophila stipulacea* has the possibility to become more prevalent in the Mediterranean basin, exacerbated by the rapid increase of water temperature. Molecular profiling appears a promising tool to study the traits that render *H. stipulacea* tolerant and resilient and facilitate its rapid and vast geographical spread. Taking advantage from recent seagrass genomes sequencing, proteomics specialty has been applied to several seagrasses giving new insight on the biology and physiology of this group of angiosperms. Thus, it could be of interest to apply proteomics to *H. stipulacea* that it could be considered as a possible plant model species to study marine biological invasion. The first step to achieve this goal is to obtain high quality proteins from plant tissue. Tissue fixation and protein extraction protocol are the most challenging steps in proteomics. Here we report a fine-tuned procedure obtained by comparing protein yield from *H. stipulacea* plants frozen in liquid nitrogen or preserved in *RNAlater* and processed following two different extraction protocols. Higher protein yield have been extracted from the procedure that use the RNA*later* preserved plants, extracted with trichloroacetic acid in water followed by trichloroacetic acid in acetone, compared to those obtained from all other procedures. Protein purity of these samples have been tested by the separation in SDS-PAGE comfirming a better resolved profile of peptide bands suitable for a gel-based proteomics. Then, to assess the quality of proteins the *m*HPLC-ESI-MS/MS mass spectrometry analyses and bioinformatics have been performed. Hundreds proteins have been identified against several seagrass genomic resources available at UniProt, NCBI, SeagrassDB and transcriptomic datasets, which were merged to form the first customized dataset useful for *H. stipulacea* proteomic investigations.

## Introduction

*Halophila stipulacea* (Forsskål) Ascherson (1867) is a native seagrass species of the Red Sea and the Indian Ocean. It entered the Mediterranean Sea in 1869 following the opening of Suez Canal and it was first recorded in the south-east Greece (Rhodes island) in 1894 (Boudouresque et al., 2009). Currently the species expands in the Eastern and Central Mediterranean Sea until Tunisia, but its occurrence is predicted to expand all over the Mediterranean Sea over this century (Georgiou et al., 2016), which might have implications for the balance between *H. stipulacea* and its native counterparts.

To understand the traits that render this seagrass tolerant and resilient toward the environmental constrains, methods and strategies to apply the molecular specialties to *H. stipulacea* have be launched (Procaccini et al., 1999; Nguyen et al., 2020a; Nguyen et al., 2020b; Winters et al., 2020). Very recently, the first draft whole-genome assembly of a *H. stipulacea* has been built (Tsakogiannis et al., 2020) whose complete validation and annotation have been expected to be released soon, so that, in coming years, gene expression studies through transcriptomics and proteomics are expected to increase. Many are the advantages offer by the proteomics specialty in marine environments; it is possible to assign function to proteins and elucidate the related metabolism in which the proteins act under different environments, e.g. in polluted or pristine areas (Nunn and Timperman, 2007; Johnson and Browman, 2007; Serra and Mazzuca, 2011); proteomics also provides a comprehensive insight into the protein profile of an organism thus revealling changes in gene expression in a complementary way to transcriptomics, as transcripts are generally loosely correlated with their corresponding proteins (Schwanhäusser et al., 2011). Finaly, the quantitative protein-level measurements of gene expression characterize biological processes and deduce the mechanisms of gene expression control and allows researchers to obtain a quantitative description of protein expression and its changes under the influence of biological perturbations, the occurrence of post-translational modifications and the distribution of specific proteins within cells (Anderson and Anderson 1998).

For all these advantages, proteomics have been applied successfully to seagrass research and have contributed in the elucidation of the metabolism dynamics and the seagrass photophysiology in the plants acclimation along a depth gradient of the ecological relevant species *Posidonia oceanica* (Procaccini et al., 2017; Mazzuca et al., 2009; Dattolo et al., 2013); proteomic approach has also revealed the behavior of the light stress-response reprogramming in the *Zostera muelleri* (Kumar et al., 2017); trough proteomics it has been elucidated the metabolic changes of the euraline *Cymodocea nodosa* in the response to manipulated salt concentrations in mesocosm (Piro et al., 2015) and give inside on the mechanisms of the adaptation to the sea acidification in natural populations of *C. nodosa* living close to volcanic CO_2_ vents (Piro et al., 2020).

When it comes to *H. stipulacea*, proteomic approach might contribute to resolve the complexity of the plant and its environment and plant-to-plant interactions during invasion by providing novel insights into cellular and biochemical pathways under contrasting conditions and contributing to identification of protein biomarkers that characterize such non-indigenous species. This might contribute to render this plant a model species to study the biological invasion of the Mediterranean sea thus justifying the efforts in developing methods to apply molecular tools. Applying proteomics to seagrass, in fact, leads two main challenges to be considered, the plant fixation method after sampling and the extreme difficulty in obtaining highly purified protein samples from tissue.

Regarding tissue fixation, seagrass samples for proteomics are usually frozen in liquid nitrogen (Mazzuca et al., 2013); this is very often uncomfortable due to the logistics of sampling, such as field experiments in remote sea regions and areas with insufficient infrastructure to allow for access to liquid nitrogen necessitate the use of fixative. So far, no other fixation method has been tested to be compared with the cryopreservation on the seagrass sample quality for proteins purification. As reported in Kruse et al., 2017 the use of the RNA*later* solution is the reliable alternative to snap freezing samples for transcriptomics and proteomics studies in plants; then, the present study aims to test the effects of the RNA*later* fixation on protein extraction efficiency and quality from *H. stipulacea* in comparison with protein yield from liquid nitrogen frozen tissues.

Quality of protein extraction for proteomic analyses significantly impacts the downstream capability of the mass spectrometers to detect peptides and the efficiency of their identification. The tissue conditions strongly influence the extraction process and then protein quality (Spadafora et al., 2008; Wang et al., 2006). The conditions of marine plant tissues, that include low protein concentration, salt enrichment, as well as compounds such as polysaccharides, lipids, phenols and other secondary metabolites, usually interfere with plant proteins separation and analyses. Moreover, biochemical conditions in seagrass tissues are extremely species specific and strongly influenced by external stress (Zidorn, 2016); for this reason, a standardized protocol for seagrass protein extraction and purification doesn’t work.

Several protein extraction protocols, in fact, have been refined to produce well-resolved electrophoretic patterns in seagrasses (Piro et al., 2015; Dattolo et al., 2013; Mazzuca et al., 2009; Migliore et al., 2007; Spadafora et al., 2008; Jiang et al., 2017); all these reports reinforce the idea that each seagrass species require an own optimized procedure to extract high-quality protein samples for proteomic approach. *H. stipulacea* has uniqueness in own metabolites repertoire, yielded two structurally macrocyclic diterpene glycoside methylglucaryl derivatives (Gavagnin et al., 2007; Carbone et al., 2008), moreover flavonoids apigenin, genkwanin and chrysoeriol (Mollo et al., 2008) that have been discussed as the molecular bio-invasion effectors (Mollo et al., 2015). On these bases, the main aim of this work is to optimize a fine-tuned procedure for *H. stipulacea* to obtain the maximum yield and quality of proteins *i)* starting from samples of *H. stipulacea* preserved in the *RNAlater* solution or frozen in liquid nitrogen; *ii*) comparing the protein yield from both preserved samples using a previous protocol applied to the iconic seagrass *Posidonia oceanica* (Spadafora et al., 2008) and the new extraction protocol developed in this study; *iii*) assessing the protein quality from two protocols by means of the SDS-PAGE; *iv*) applying the gel-based proteomics coupled with mass spectrometry analyses to the sample of proteins showing higher purity and quantity.

As a reference genome for *H. stipulacea* does not exist yet, the protein identification will be made using a customized database built with sequences from the complete genomes of two seagrass species, *Zostera muelleri* and *Zostera marina* coupled with transcriptomic datasets from *Cymodocea serrulata, Halophila ovalis, Posidonia oceanica* downloaded from several database repositories.

## Materials and methods

### Sample preparation

*H. stipulacea* plants have been collected by SCUBA diving from meadows expanding the island of Crete, Greece (Maridati 35.22183°N 26.27310°E in summer 2018 and Atsikari 35.255°N, 26.2233° E in summer 2019). The samples (individuals or *genets*) each formed by a rhizome and three to four shoots (*ramets*) were cleaned from epiphytes, washed rapidly in water and frozen in liquid N_2_ or fixed in *RNAlater* following the manufacturer’s instructions (ThermoFisher Scientific, Waltham, Massachusetts, US); in short, plants were immerged in *RNAlater* in small vials, kept at 4 ° C for few days and then have been stored at - 20°C. N_2_ frozen samples were kept at -80°C. Under these conditions both kind of samples have been stable for several months till the protein extraction.

### Extraction and purification of total protein from Halophila

Protein extraction has been performed after three-five months from the samples harvesting. Two procedures were applied for protein extraction and purification from *H. stipulacea* tissues, the Procedure 1 optimized in this work and the Procedure 2 developed for *P. oceanica* by Spadafora et al., 2008. Procedures differ in the amount of tissue used for the protein extraction and in chemicals that were used in the step for removal of interfering molecules from tissues prior to purify proteins by the phenol phase. Details of both procedures are reported in the Figure 1. A reciprocal approach has been also applied: samples in *RNAlater* have been extracted with Procedure 2 and samples fixed in liquid nitrogen have been processed with Procedure 1.

**Figure 1.**
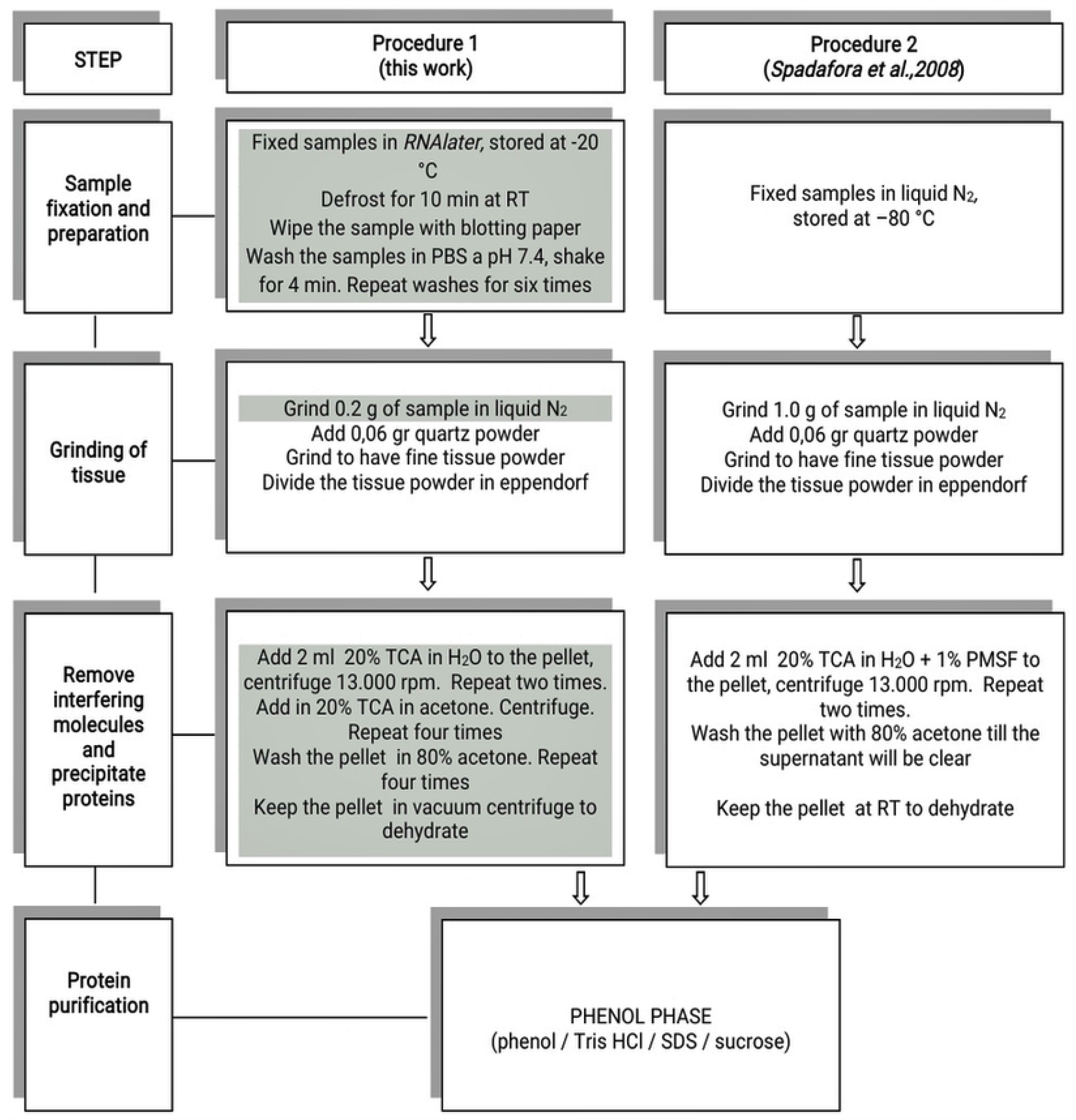
Details of two extraction procedures and comparison among the various steps applied to extract proteins from *Halophila stipulacea* tissue. The steps that differ between two procedures are marked in gray.

For protein purification approximately 0.1g of powdered tissue from each procedure was dissolved in 0.8 ml of phenol (buffered with Tris HCL, pH8.0, Sigma, St. Louis, MO, USA) and 0.8 ml of SDS buffer (30% sucrose, 2% SDS, 0.1 M Tris-HCl, pH 8.0, 5% 2-mercaptoetanol) in a 2 ml microfuge tube. The samples were vortexed for 30 s and centrifuged at 13000 rpm for 5 min to allow proteins to melt in the phenol phase. The phenol phase was mixed with five volumes of 0.1 M ammonium acetate in cold methanol, and the mixture was stored at − 20°C for 30 min to precipitate proteins. Proteins were collected by centrifugation at 13000 rpm for 5 min. Two washes were performed with 0.1 M ammonium acetate in cold methanol, and two with cold 80% acetone, and centrifuged at 13000 rpm for 7 min. The final pellet containing purified protein was dried and dissolved in Laemmli 1DE separation buffer overnight (Laemmli, 1970). Proteins were then quantified by measuring the absorbance at 595 nm according to the Bradford assay. Protein yield was calculated as mg of protein for g fresh tissue weight in three biological replicates for each sample. For each replicate two independent extractions have been made. The relative abundances of proteins were calculated as a mean value ± standard error (n = 6). A Student t-test was used to make pair-wise comparisons between samples. Unless otherwise noted, p-levels of 0.05 were used as the threshold for statistical significance.

### Electrophoresis of leaf proteins, protein in-gel digestion and mass spectrometry analyses

Gels preparation and electrophoreses of samples were carried out according to the method of Laemmli, 1970. The ratio of acrylamide/bisacrylamide was 12.5 % in the running gel and 6 % in the stacking gel. The samples were heated for 5 min at 100 °C before being loaded on the gel at the amount of 5, 10 and 20 µg for both extraction protocols. The electrophoretic run was carried out in running buffer at 60 mA for the stacking gel and 120 mA in the running gel at constant power of 200 V, for 1 h and 15 min. The gels were stained with Coomassie Blue overnight and subsequently destained with several changes in the destaining solution (45% methanol, 10% acetic acid). Digitalized images of the destained SDS-PAGEs were analyzed by the Quantity One 1-D Analysis Software (Bio-Rad, Berkeley, US) to measure the band densities at each lane of all biological replicates; the amount of protein at bands of 55, 25, and 10 kDa was done using the marker reference bands at 75, 50, and 25 kDa that contained 150, 750, and 750 ng of proteins respectively (Figure 2). Each lane of the same SDS-PAGE was divided in six slices from 200 to10 kDa and manually excised from the gel.

**Figure 2.**
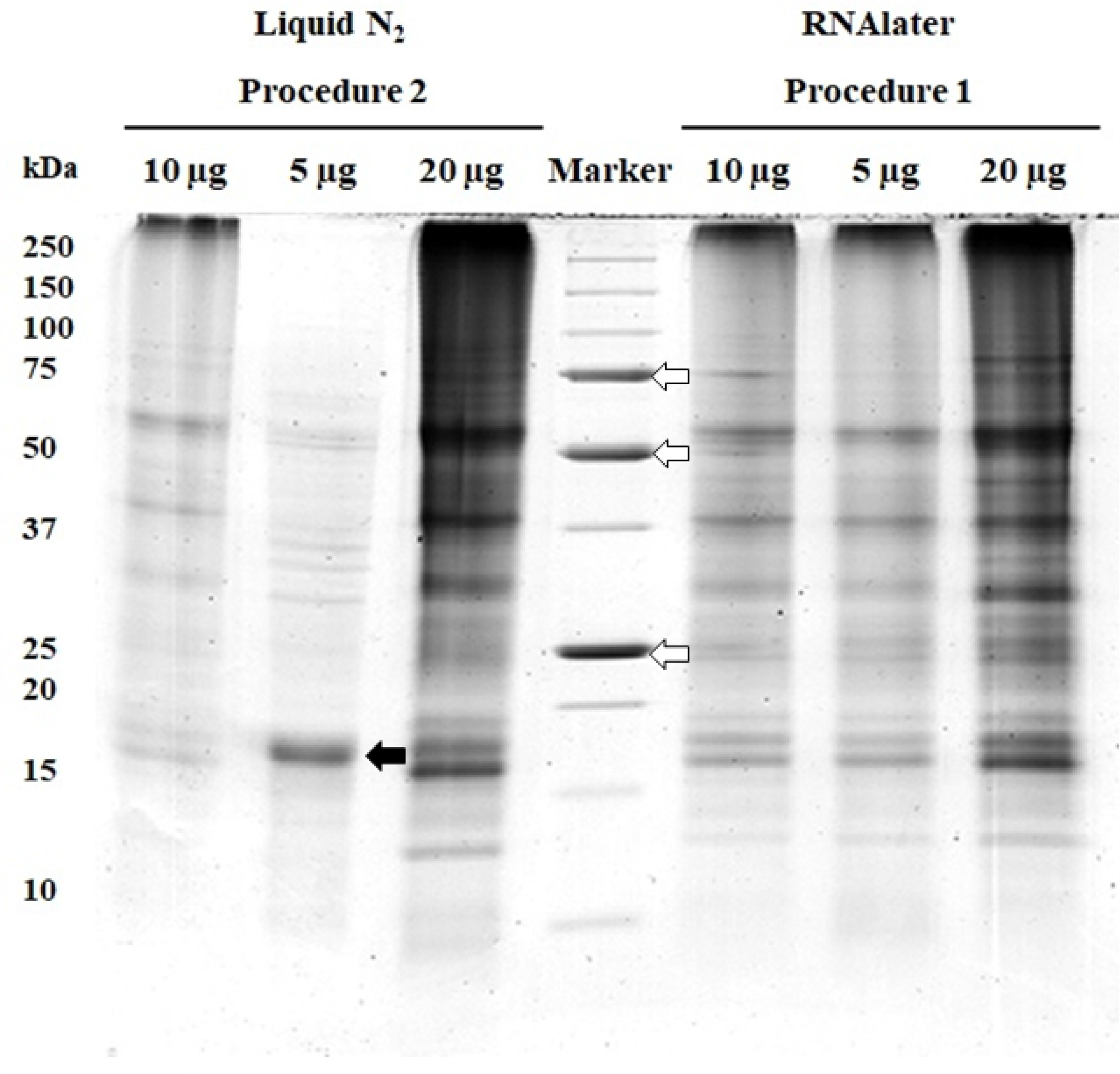
SDS-PAGEs of purified proteins extracted from *Halophila stipulacea* plants frozen in liquid N_2_ and following the Procedure 2 or fixed in RNAlater and following the Procedure 1. Samples were loaded at different amount of 5, 10 and 20 µg for both extraction protocols. The white arrows indicate the marker bands that have been quantized by means of the Quantity One software. The black arrow indicates the major polypeptide appeared in the lane loaded with 5 µg proteins from the N_2_-Procedure 2 (see details in the text).

The CBB-stained gel slices were destained in 50 mM ammonium bicarbonate and acetonitrile (ACN) (1:1 v/v) and then processed with reduction/alkylation steps with DTT at 56°C, 20 min and 55 mM iodacetammide at RT, 30 min in the dark (Shevchenko et al., 2007). Reduced and alkylated gel pieces were processed for in-gel protein digestion by trypsin (Promega, Madison WI, USA) overnight at 37 °C adding ammonium bicarbonate buffer to cover gel matrix. The tryptic peptides were extracted with 5% formic acid (FA) in water and then washed in acetonitrile and ammonium bicarbonate (50 mM). Samples were dried and dissolved in 20 µl of 8% formic acid in water.

### Tandem MS analysis

Twenty microliters of tryptic digested peptides were injected on a reversed phase trap column (Analytical Column LC18 BioBasicTM, 300 Å, 5 µm, 50 µm ID × 1 mm length, Thermo Scientific, US). Separations were performed using an ultra-chromatographic system (UltiMate 3000 RSLC System, Thermo Scientific, US) at a constant flow rate of 100 µL/min with a gradient from 4% buffer A (2% ACN and 0.1% FA in water) to 96% buffer B (2% water and 0.1% FA in ACN) in 60 minutes. The eluting peptides were on-line sprayed in a LTQ XL mass spectrometer (Thermo Scientific, Sacramento, US). Full scan mass spectra were collected in the linear ion trap in the mass range of m/z 350 to m/z 1800 Da and the 10 most intense precursor ions were selected for collision-induced fragmentation. The acquired MS spectra were used for protein identification.

### Bioinformatics analysis and Peptide identification of proteins of Halophila

#### Local database

A customized local database for protein identification and functional annotation was built using the FASTA deduced sequences from i) *Zostera marina* genomes and *P. oceanica* transcriptomic sequences from NCBI and UniProt (downloaded in February 2020), ii) customized peptide dataset from *Posidonia oceanica* transcriptomes (Dattolo et al., 2013), iii) peptide dataset from of *C. serrulata* and *H. ovalis* transcriptomes, stored at the SeagrassDB (Sablok et al., 2018).

The spectra in raw format were interfaced with the local database using the PatternLab for Proteomics software and converted into the .*sqt* format, useful for identification using Scaffold (version Scaffold_4.11.0, Proteome Software Inc., Portland, OR) (Carvalho et al., 2016).

Scaffold was used to validate MS/MS based peptide and protein identifications. Peptide identifications were accepted if they could be established at greater than 95.0% probability by the Peptide Prophet algorithm (Keller et al., 2002) with Scaffold delta-mass correction. Protein identifications were accepted if they could be established at greater than 99.9% probability and contained at least 2 identified peptides. Protein probabilities were assigned by the Protein Prophet algorithm (Nesvizhskii et al., 2003).

#### Database Searching

All MS/MS samples were analyzed using Sequest (Thermo Fisher Scientific, San Jose, CA, USA; version N/A). Sequest was set up to search the local customized database assuming carbamidomethylation of cysteine as fixed modification and the digestion enzyme trypsin. Sequest was searched with a fragment ion mass tolerance of 1.0 Da and a parent ion tolerance of 40 ppm.

Auto MS/MS spectra were extracted from raw data accepting a minimum sequence length of ten aminoacid and merging scans with the same precursor within a mass window of ±0.4 m/z, in a time frame of ±30 s.

Auto thresholds were used for peptide identification in Scaffold software. Generally, peptide probabilities are assessed using a Bayesian approach to local FDR (LFDR) estimation to achieve a target of 2.35%. Functional annotations of further unidentified sequences have been made by the OmicsBox Base Platform (BioBam Bioinformatics S.L., Valencia, Spain) against the NCBI Viridiplantae database downloaded on October 9, 2020.

Second level GO categories of biological process, molecular function, and cellular components among the annotate protein of *H. stipulacea* were obtained with BLASTP tool available at UniProt database.

## Results

### Protein extraction and protein yield of H. stipulacea samples

Tissues of *Halophila* gave a different average total proteins yield depending on the sample fixation and extraction procedures, as reported in Table 1. The protein yield from *RNAlater* samples and extracted with the Procedure 1 was higher than protein yield of samples fixed in N_2_ and extracted following the Procedure 2; in our conditions, no significant difference in protein yield was observed among *genets* and *ramets* belonging at the same set of samples. The reciprocal approach among tissue fixation methods and extraction procedures, gave lower protein yield in both cases; this suggest that the removal of interfering molecules/protein precipitation is the key step that is affected by the tissue fixative as well as by the chemicals used in two procedures (Table 1).

**Table 1.**
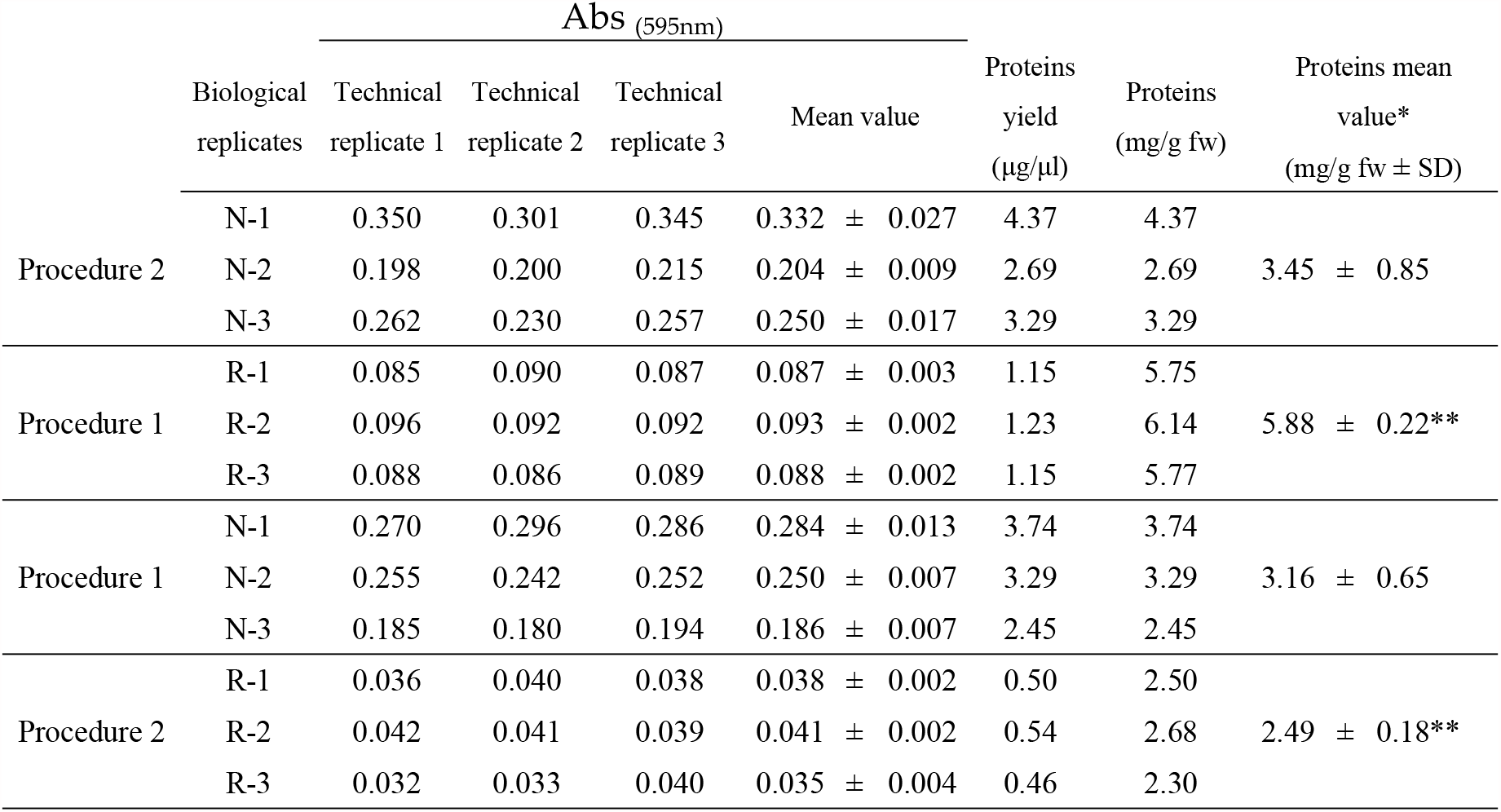
Spectrophotometrical absorbance (Abs) and purified protein yields obtained from *Halophila stipulacea* plants frozen in liquid nitrogen (N) or fixed in *RNAlater* (R). *Values are the mean of three biological replicates and three technical replicates (P<0.05); ** protein extraction was made from 200 mg fresh tissue

As can be seen in Figure 2, the SDS-PAGE of proteins from the N2 fixed plants and purified according to the Procedures 2 generates a non linear increase in number and intensities of the polypeptide patterns in the 20, 10 and 5 µg lanes; in this last, a prominent polypeptide band appeared, with the apparent molecular weight of 18 kDa that is not resolved in the 10 µg and 20 µg lanes; conversely, the 20 µg lane shows polypeptide bands that are more than twice as strong as those in the 10 µg lane. Take all together these findings are consistent with the persistence of residual contaminants in samples, that interfere with the denaturation by heating in presence of thiol reagents and excess of SDS. In comparison, proteins from the RNA*later* fixed plants, purified with the Procedure 1, gave well resolved number of sharp bands of polypeptides without background in all lanes; moreover, intensity and number of bands increase accordingly with protein amount loaded in each lane, indicating the purest quality of proteins. The patterns of proteins from *RNAlater*, have prominent bands at 55 kDa, 37 kDa, 30 kDa and 15 kDa.

Protein samples coming from the reciprocal approach showed lesser resolved and lesser abundant bands with intense vertical streaking in the SDS-PAGE lanes in both cases, thus suggesting that the reciprocal approach do not remove efficiently the interfering compounds, affecting the protein quality besides the protein yield (Figure 3; Table 1). As the better SDS-peptides profiles the better is the protein quality and purification, the mass spectrometry analyses have been addressed only at protein samples coming from the samples fixed in *RNAlater* and processed with the Procedure 1.

**Figure 3.**
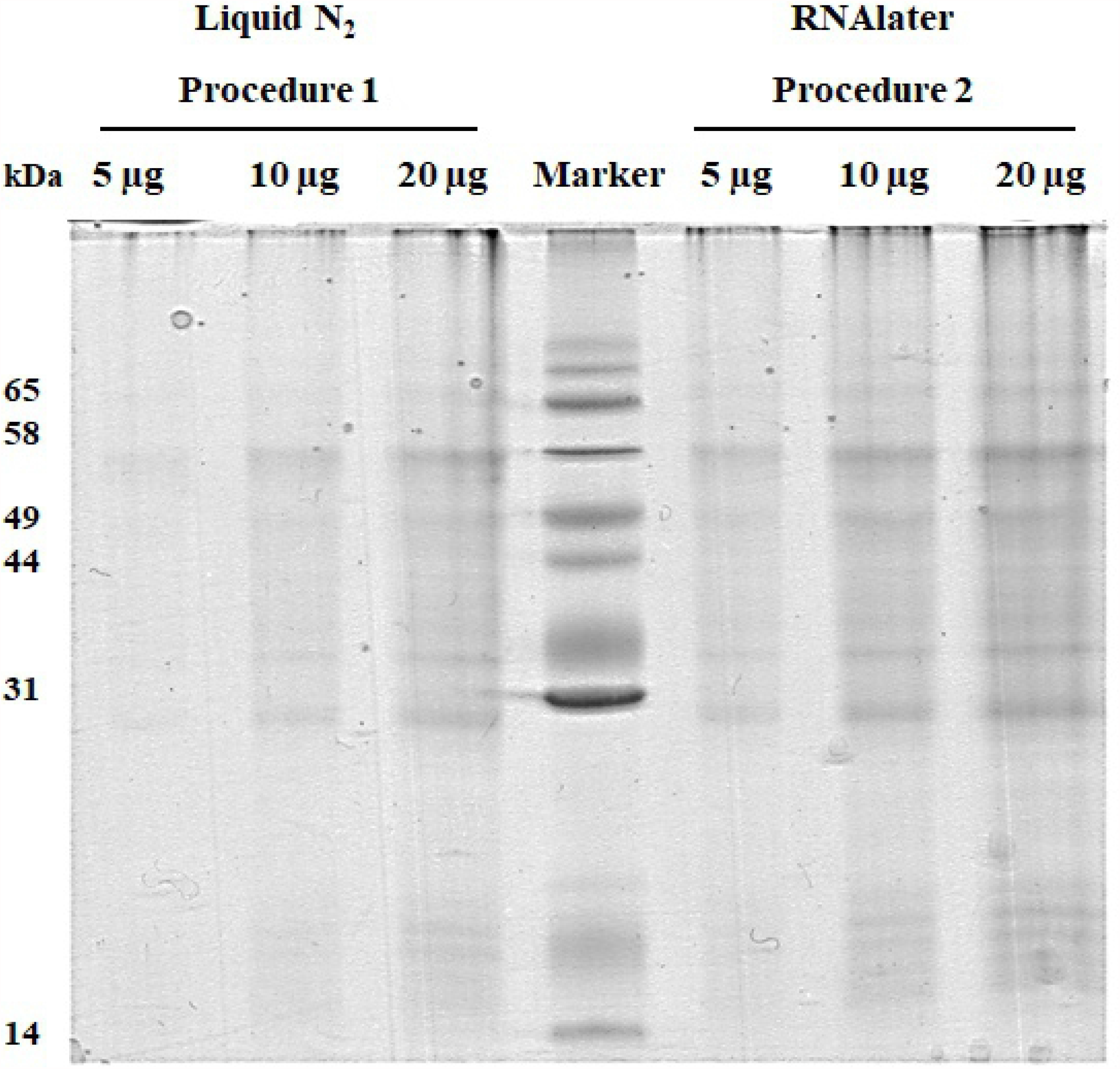
SDS-PAGEs of purified proteins extracted from *Halophila stipulacea* plants frozen in liquid N_2_ and following the Procedure 1, fixed in *RNAlater* and following the Procedure 2. Samples were loaded at different amount of 5, 10 and 20 µg for both extraction protocol.

### H. stipulacea protein identification against genomic and transcriptomic seagrass databases

Functional annotations of the identified sequences have been performed, first, against the generalistic UNIPROTKB_VIRIDIPLANTAE and NCBI *Viridiplantae* protein databases excluding *Z. marin*a, *Z. mueller*i and *P. oceanica* sequences from the analysis; this in order to identify as many proteins as possible using a larger sequence database, but also to measure the gap of the sequence homologies of *H. stipulacea* toward the terrestrial plants and among the unrelated species to seagrasses. The enquiring gave more than 5,000 functional annotations (Supplementary Table 1), but only eighteen had two spectra for each protein to satisfy the minimum required for a significance of protein identification score (Eriksson et al., 2000); the rest of the identifications, had only one spectrum for each protein and then they have been not considered in this study (Supplementary Table 2).

By using the customized local database, 889 functional annotations have been identified, each with not less than 2 peptides per proteins, not less than two spectra each peptide and not less than 94% peptide identification probability. Peptide sequences, statistical parameters obtained from the alignments of all identified proteins in all analyzed samples are reported in the Supplementary Table 2 and 4.

The bar plot in the Figure 4 shows the number of *H. stipulacea* proteins identified with each database and species. The *Z. marina* genome from UniProt and NCBI repositories gave 144 and 136 functional annotations respectively; dataset from *P. oceanica* available at the NCBI repository, gave no significant annotations; 31 protein sequences were identified from the customized dataset from *P. oceanica* transcriptomics. Identification from the *H. ovalis* transcriptome dataset gave 141 identified proteins, and 167 proteins were recognized against the *C. serrulata* dataset.

**Figure 4.**
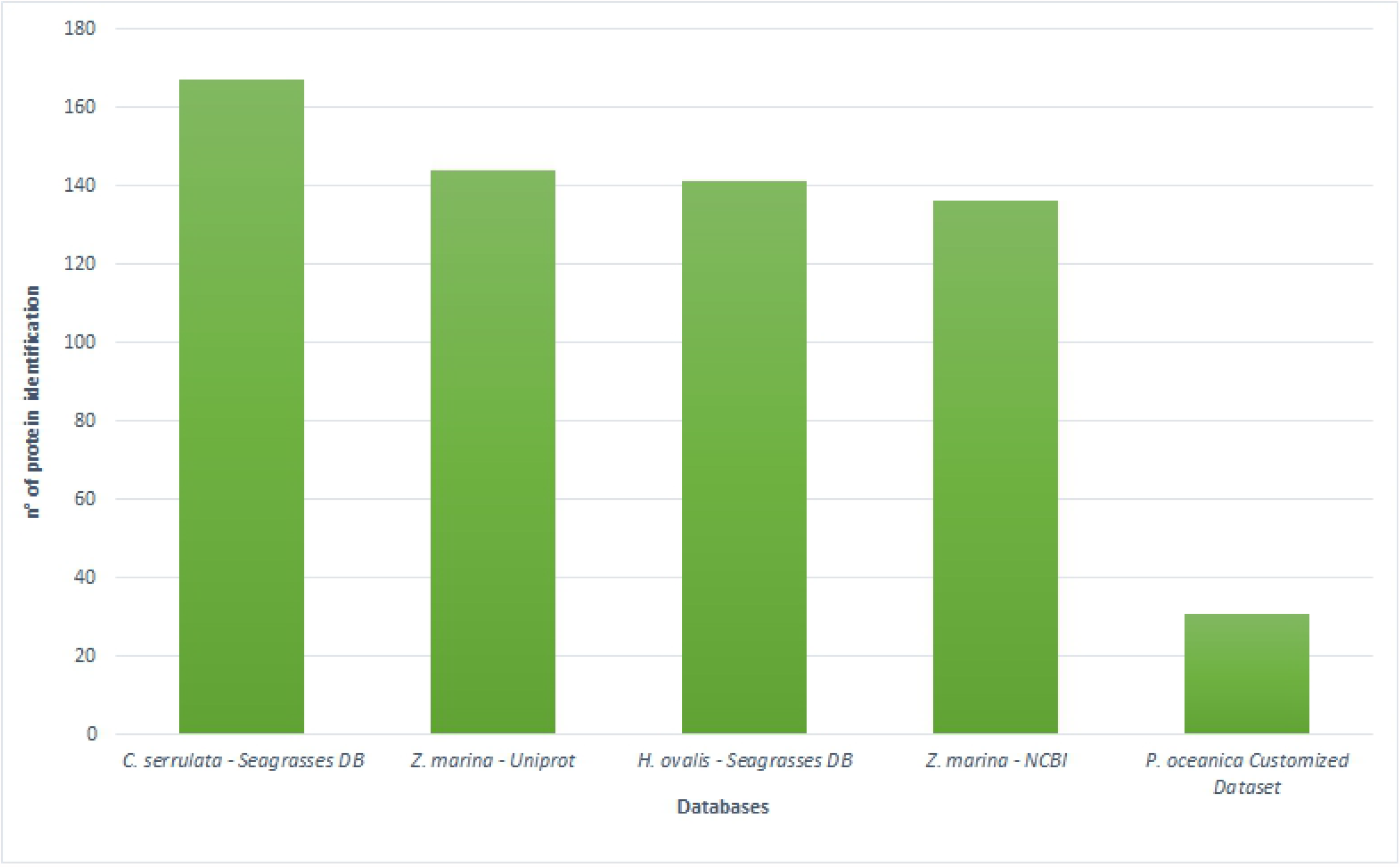
Bar plots show the number of identified proteins of *H. stipulacea* obtained against genomic and transcriptomic sequence datasets from four seagrass species, available at public repositories and customized resources.

Interestingly, 270 identifications have been found belonging sequences of the bacteria *Planctomycetes bacterium* KOR34 strains, recently renamed as *Posidoniimonas corsicana* (Kohn et al., 2020) and *Marinomonas posidonica* IVIA-Po-181 strain (see all the identified proteins in the Supplementary Table 4).

In the Figure 5 the Gene Ontology analyses, made by the UniProt tool interrogated on October 30, 2020, shows that among the category “molecular function”, “cellular component” and “biological processes”, most proteins are belonging to the sub-categories “binding”, “catalytic activity”, “cellular anatomical entity”, “cellular process”, “metabolic process”, “biological regulation”. Details of GO assignment are reported in the Supplementary Table 3.

**Figure 5.**
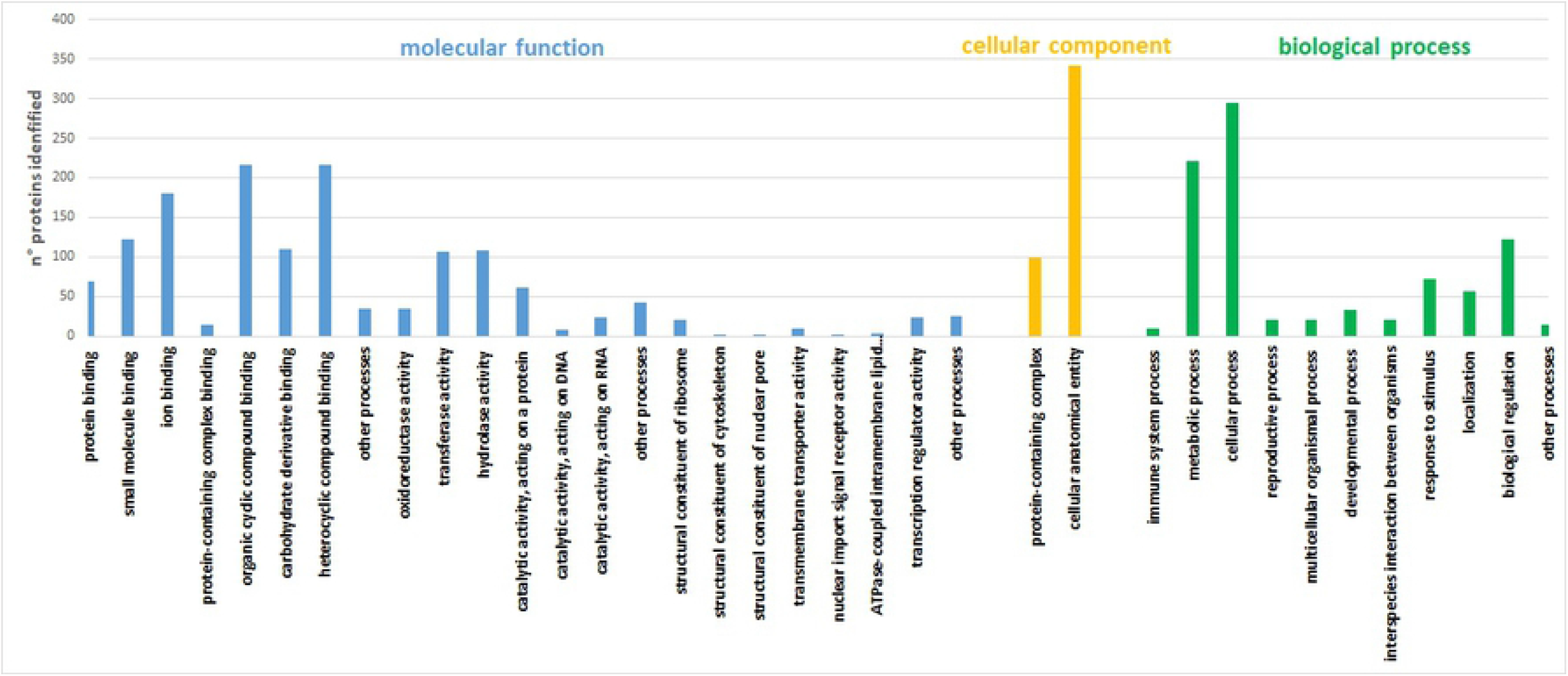
Number of identified proteins of *Halophila stipulacea* belonging to the categories of Gene Ontology, analyzed by Uniprot database tool (in October, 2020).

The biological functions, with large number of identified proteins, are the cytoskeleton metabolism whose Actin-related protein, Microtubule-associated proteins, formin-like protein, protein-tyrosine-phosphatase MKP1-like and various myosin isoforms deputated to the vesicle transport along actin filaments have been detected; many proteins belonging to the carbohydrate metabolic process such as Transaldolase, Polygalacturonase, Fructose 2,6-bisphosphate. Cell wall organization and cell wall biogenesis are well represented with Kinesin-like protein, Hexosyltransferase involved in the pectin biosynthesis, Endo-1,4-beta glucanase, Cellulose synthase-like CSLD, Glucomannan synthase and many others involved in the cellulose metabolism. Chloroplast biological functions count 35 identified proteins such as Outer envelope protein 80, K(+) efflux antiporter 2, Bifunctional aspartokinase/homoserine dehydrogenase 2, Violaxanthin de-epoxidase, Protein ACTIVITY OF BC1 COMPLEX KINASE 1 responsive to nitrogen starvation. Defense response gather 11 proteins, the Ethylene-responsive transcription factor 1, disease resistance protein RGA2, Protein kinase 1A, Glutathione S-transferase F6 and others. DNA and mRNA metabolisms count more than 30 identified proteins, including replication, repair, recombination, transcription, and splicing. Large identifications are belonging to the anabolic and catabolic metabolisms of proteins, PTM and transport. All detailed results are reported in the Supplementary Table 3.

## Discussion

Here we provide a fine-tuned method to apply the molecular technologies for protein expression analysis of *H. stipulacea* take advantage from the seagrass genomic resources available so far and from its own genome once it will be fully annotated. By applying the well-established protocol developed for *P. oceanica*, in fact, a lower protein yield and poorer protein quality from *H. stipulacea* tissue have been obtained in comparison with those obtained with the protocol developed in this study, demonstrating that the specific chemical moieties of *H. stipulacea* tissue, that differs from other seagrasses, imposes new procedure for extracting high quality proteins. The extraction procedure optimized in this work strengthens the protein precipitation step by the trichloroacetic acid in water, thereby improving the removal of water-soluble interfering compounds, followed by a further protein precipitation by trichloroacetic acid in acetone that removed the water-insoluble molecules more efficiently than the compared protocol.

The new protocol also uses five-times lower tissue amount than those from other procedures applied in seagrasses (Piro et al., 2015; Mazzuca et al., 2009) and use, for the first time, the fixation with *RNAlater*, instead than liquid nitrogen, that favors the yield and the quality of the extracted proteins. Plant fixation alternative to freezing, might be easily performed in unsuitable places such as boats, harbors, or place very far from the equipped laboratories, thus facilitating the sampling and shipping of plants for molecular analysis. The *RNAlater* fixation, in fact, could reduce the times elapsed between the sampling at sea and the freezing of the samples at lab, which are important in comparative proteomic studies (Mazzuca et al., 2013).

Botton-up proteomic approach and gel-based mass spectrometry have been applied for the first time to *H. stipulacea*. We were able to identify proteins that found their significant sequence homology against sequence datasets from the seagrasses *H. ovalis*, a directly related species to *H. stipulacea, C. serrulata, Z. marina* and *P. oceanica*; identifications were merged to form a customized dataset useful for *H. stipulacea* proteomic investigations. The dataset might be implemented by sequences coming from the genome sequencing of the species and that, at moment, are still on the way to be fully validated and annotated (Tsakogiannis et al., 2020).

The wider identification of proteins was obtained against the genome sequence database of *Z. marina* available at the NCBI repository and UniProt database; minor identifications were made against *C. serrulata* and *H. ovalis* at the SeagrassDB; the reason why is that NCBI and UniProt have several well annotated sequences coming from a complete genome sequencing. Regarding the genetic categories in which the identified proteins fall, the categories linked to the DNA and RNA metabolisms, the protein synthesis and degradation via ubiquitination, the cell wall and cytoskeleton metabolisms, the defense responses toward biotic and abiotic stress are well represented; surprisingly the photosynthetic metabolism, generally represented by proteins belonging to the membrane-bound photoreceptor complexes PSI and PSII, is not well represented. A further rather unusual finding in a plant proteomic analysis was the lack of identification, using the NCBI database and SeagrassDB, of the enzyme Ribulose bisphosphate carboxylase (Rubisco) which is undoubtedly the most abundant protein in leaves; a possible explanation lies that since whole plants were used, the amount of leaf tissue was lower than rhizomes and roots and this could affect the final concentration of the enzyme. A further hypothesis is that sequences of Rubisco retrieved from the NCBI and SeagrassDB databases did not match with MS/MS spectra obtained from the enzyme of *H. stipulacea*. On the other hand, the Rubisco large subunit (LSU) of *H. stipulacea* (H6TQS9) at the UniProtKB/Swiss-Prot database at the ExPASy Bioinformatics Resource Portal (Aritmo et al., 2012) consists just of a fragment of 200 amino acids, which represent less than half of the total 476 residues of a typical LSU sequence. The LSU short fragment did not allow us to perform efficiently the in-silico generation of theoretical spectra to be used in the proteomic fingerprint with the experimental spectra obtained from *H. stipulacea* (data not shown). A mention apart deserves our findings on cell wall proteins, highlighting that *H. stipulacea* has many sequence homologies with the other seagrass orthologous proteins belonging to the cell wall metabolism and function. Seagrasses possess a specific cell wall structure and an exclusive repertoire of carbohydrate composition (Olsen et al., 2016), thus a specific cell wall proteome is also expected.

Bioinformatics gave also significant identifications against sequences from *Planctomycetes bacterium* KOR34 and *Marinomonas posidonica* IVIA-Po-181 that have been found associated to *P. oceanica* leaves (Kohn et al., 2020; Lucas et al., 2012) and *H. stipulacea* tissues (Weidner et al., 2020). Genomes of both bacteria strains have been sequenced, validated and annotated in NCBI and UniProt repositories.

This finding suggests that *RNAlater* fixative might also preserve mRNAs and proteins of the seagrass-associated microbiomes; in favour to this hypothesis, it has reported that *RNAlater* better preserves the marine microbial proteome in environmental sample collection in comparison with other fixatives (Saito et al., 2011). Given the relevance of the microbiome in the ecosystem services of seagrasses, this aspect may deserve further investigation.

As last result, a high number of validated peptide spectra obtained in this work have received not significant matching or peptides remained still unidentified. Low statistical significance in the identification of proteins found against the generalist UNIPROTKB_VIRIDIPLANTAE and NCBI *Viridiplantae* protein databases showed the poor correspondence in the sequence homology among the well annotated genomic resources of many terrestrial plant species and *H. stipulacea*, thus indicating a low functional and evolutionary relationships between sequences.

The availability of genomic and/or transcriptomic sequences from *Halophila* spp. could certainly reduce this gap of knolwdge by reducing the number of the orphan peptides and thus determine, in near future, a more comprehensive analysis of metabolic pathways at the level of protein expression in natural populations. In any case, this analysis definitively opens the scenario for the applications of molecular methodologies also in *Halophila* spp., like what has been done for other seagrasses.

## Conclusions

The *Halophila stipulacea* proteome dynamics might contribute to elucidate the complexity of the plant and its environment and plant-to-plant interactions with native species during the invasion of the Mediterranean basin. Thus, it is of interest to develop sound method and procedure to obtain good protein samples to be used in the proteomic pipeline. In this work we demonstrated that the chemical moieties of *H. stipulacea* tissue, that differs from other seagrasses, imposes a new procedure for extracting high quality proteins. By applying the fine-tuned procedure developed in this work, in fact, we obtain higher protein yield and quality of *H. stipulacea* plants in comparison with those obtained using a protocol optimized for *Posidonia oceanica*. This fine-tuned procedure starts from RNA*later* fixed tissue and uses, as the first step, the aqueous trichloacetic acid solution that removes water soluble interfering compounds more efficiently than the proposed compared protocol; additionally the SDS-PAGE profiles confirmed that proteins extracted by the fine-tuned procedure are of high purity and quality. Mass spectrometry and bioinformatics gave hundreds of significant protein identifications whose number depends on the seagrass database used, and more relevant, gave no significant identification against generalist protein databases, thus indicating a low functional and evolutionary relationships between *H. stipulacea* and many terrestrial plants. We expect that, once the genome of this plant will be validated and available, the procedure here developed will be very useful for the application of proteomics to the molecular ecology of *H. stipulacea* in a complementary way than all the other “omics” sciences.

## Acknowledgments

This research was co-financed by Greece and the European Union (European Social Fund-ESF) through the Operational Programme ‘Human Resources Development, Education and Lifelong Learning 2014–2020’ in the context of the project ‘I-ADAPT’ (MIS 5006611). Authors sincerely thank Gaurav Sablok and Regan Hayward who provided sequences of seagrass transcriptomes stored at SeagrassDB. We thank Julius Glampedakis and Thanos Dailianis for helping during field work. The authors thank also the project PON “Sistema Integrato di Laboratori per l’Ambiente (S.I.L.A.) – PON a3_00341A for the availability of the mass spectrometer equipments.

## Author Contributions

Conceptualization, S.M. and E.A.; Methodology, A.P. and V.A.; Software, A.P.; Validation, S.M., A.P. and E.A.; Formal Analysis, A.P.; Investigation, A.P. and V.A.; Resources, S.M. and E.A; Data Curation, S.M.; Writing – Original Draft Preparation, A.P.; Writing – Review & Editing, S.M, E.A. and A.P.; Visualization, E.A.; Supervision, S.M.; Project Administration, E.A.; Funding Acquisition, E.A.

